# Risperidone-induced changes in DNA methylation from peripheral blood in first-episode schizophrenia parallel neuroimaging and cognitive phenotype

**DOI:** 10.1101/2020.03.31.018283

**Authors:** Maolin Hu, Yan Xia, Xiaofen Zong, John A. Sweeney, Jeffrey R Bishop, Yanhui Liao, Gina Giase, Bingshan Li, Leah H. Rubin, Yunpeng Wang, Zongchang Li, Ying He, Xiaogang Chen, Chunyu Liu, Chao Chen, Jinsong Tang

**Affiliations:** Department of Psychiatry, the Second Xiangya Hospital, Central South University, Changsha, Hunan, China; Center for Medical Genetics, School of Life Sciences, Central South University, Changsha, Hunan, China; Department of Psychiatry, State University of New York Upstate Medical University, Syracuse, NY; Department of Psychiatry, Renmin Hospital of Wuhan University, Wuhan, Hubei, China; Department of Psychiatry and Behavioral Neuroscience, University of Cincinnati, Cincinnati, Ohio; Department of Experimental and Clinical Pharmacology and Department of Psychiatry, University of Minnesota, Minneapolis, MN; Department of Psychiatry, Sir Run-Run Shaw Hospital, School of Medicine, Zhejiang University, Hangzhou, Zhejiang 310016, China; Department of Psychiatry, University of Illinois at Chicago, Chicago, IL; Genetics Institute, Vanderbilt University School of Medicine, Nashville, TN; Department of Neurology, Johns Hopkins University School of Medicine, Baltimore, MD; Division of Mental Health and Addiction, Oslo University Hospital, Oslo, Norway; National Clinical Research Center for Geriatric Disorders, Xiangya Hospital, Central South University, Changsha, China; Mental Health Institute, the Second Xiangya Hospital, Central South University, Changsha, Hunan 410011, China; National Clinical Research Center on Mental Disorders, Changsha, Hunan 410011, China; National Technology Institute on Mental Disorders, Changsha, Hunan 410011, China; Hunan Key Laboratory of Psychiatry and Mental Health, Changsha, Hunan 410011, China

**Keywords:** Biomarker, DNA methylation, risperidone treatment, first-episode schizophrenia

## Abstract

Today, second generation anti-psychotics such as clozapine and risperidone are the favored treatment for schizophrenia. Yet, the absence of relevant biomarkers that can decode their neurobiological effect shackles our ability to accurately predict and track response to treatment. While researchers have investigated DNA methylation as a biomarker for schizophrenia risk, none have performed a systematic analysis of the effect of antipsychotics upon DNA methylation. We hypothesize that disease-related methylation changes occur before treatment, and that acute antipsychotic treatment may affect DNA methylation. We designed a longitudinal DNA methylation study to estimate risperidone’s effect on DNA methylation and how changes in DNA methylation might influence risperidone’s therapeutic effect on behavioral and neuroimaging phenotypes. Thirty-eight patients with first-episode drug-naïve schizophrenia (FES) and 38 demographically-matched individuals (healthy controls) participated. We identified brain related pathways enriched in 8,204 FES-associated methylation sites. Risperidone administration altered methylation in 6,143 CpG DNA sites. Post-treatment FES associated with methylation in 6760 CpG sites. Majority of the DNA methylation changes were treatment effect in the overall CpG sites, the FES associated CpG sites, and risperidone associated CpG sites, except for the post-treatment FES associated CpG sites. There were 590 DNA methylation cites normalized by risperidone treatment. The methylation changes of these 590 CpG sites were related to alterations in symptom severity, spontaneous neurophysiological activity, and cognitive function. To our knowledge, this is the first longitudinal methylation study of drug treatment effect and side effect in psychiatric disorders to include parallel studies of neuroimaging and cognitive phenotypes. We identified FES-associated CpG sites not confounded by drug treatment as potential SCZ biomarkers. The normalization effect of risperidone monotherapy suggests that DNA methylation changes may serve as a predictive biomarker for treatment effect. The constructed methylation-phenotype network revealed a relationship between methylation and a wide range of biological and psychological variables.

## Introduction

For the majority of patients with schizophrenia (SCZ), treatment features second-generation antipsychotic drugs such as risperidone supported by psychosocial therapy. Risperidone has receptor affinities targeting dopamine, serotonin and other neurotransmitters.[1] Risperidone effectively treats acute psychosis and works to prevent SCZ relapse.[2] Although antipsychotic drugs bind to target receptors within hours of administration, clinical efficacy can take weeks, perhaps due to slower acting complex biological changes.[3] Accumulating evidence indicates that gene expression regulation, such as DNA methylation, may spark the clinical effect of antipsychotic drugs,.[4–6] The epigenome-wide association studies of SCZ disease risk to date, a mix of disease-discordant monozygotic twins or case-control comparisons, [7–17] produced inconsistent results regarding changes in DNA methylation.[8] These inconsistencies may result from wide variance in illness duration and drug treatment histories of the patients studied.

Previous studies have shown antipsychotics influencing DNA methylation,[18–26] with olanzapine changing DNA methylation in the brains of mice[20], and clozapine changing DNA methylation in human peripheral leukocytes.[27] Again, studies to date involved primarily chronically treated patients with long histories of disease symptoms and varied drug treatments. Under these circumstances disease severity may muddle a clear measure of treatment effect. Alternatively, studying drug-naïve patients with only a first episode of SCZ (FES) would avoid these likely confounders and produce a clearer indication of the effects of antipsychotics on DNA methylation.

We have two aims in the current study: 1) identifying disease-related methylation changes before treatment, and 2) investigating acute antipsychotic treatment effects on DNA methylation. We hypothesize that disease-related methylation changes occur before treatment, and that acute antipsychotic treatment may affect DNA methylation. We used a longitudinal cohort design to study the effects of the risperidone monotherapy on DNA methylation in patients with FES (Fig.1). We followed the drug naïve patients over the treatment course, recording phenotypic changes and monitoring DNA methylation. We also integrated retrospectively collected data from demographic matched controls to help differentiate treatment effect from the side-effect of methylation changes. We then explored whether risperidone-induced methylation changes correlated to brain phenotypes, with features including symptom severity, cognitive function, and spontaneous brain activity.

**Fig.1.**
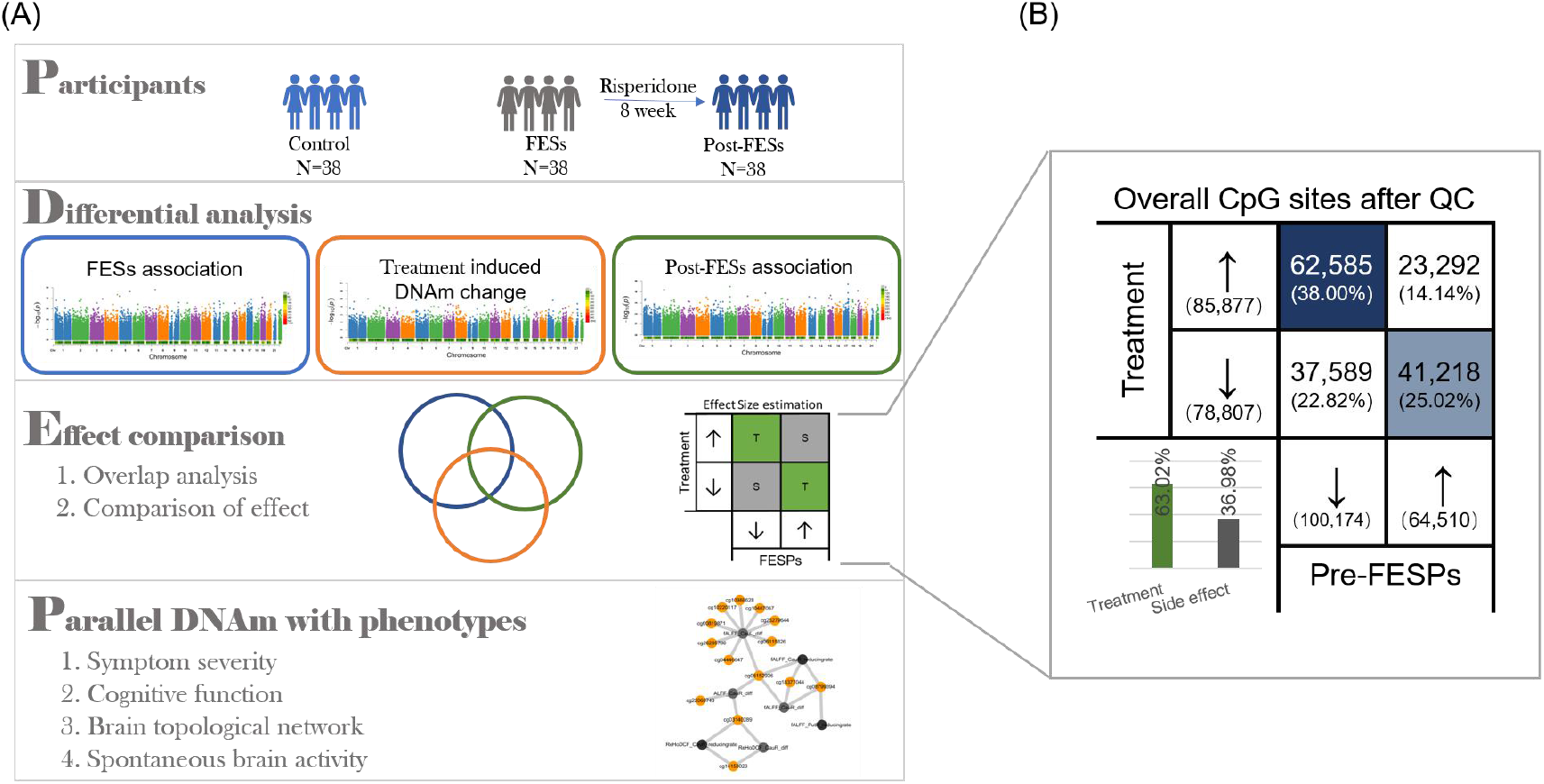
Overall study design. (A)1. Thirty-eight patients with first-episode drug-naïve schizophrenia (FES) and 38 demographically-matched individuals (healthy controls) participated. 2. We conducted differential analysis among FESs vs Control, post-FESs vs FESs, and post-FESs vs Control. 3. Comparison of the differential results. 4. Parallel DNA methylation changes with phenotypes changes. (B) Overall comparison of effect size between two differential analysis: pre-treatment FEP association analysis (pre-FEP vs control) and treatment association analysis (Post-FEP vs Pre-FEP)

## Results and Discussion

### 1. FESs associated methylation enriched in neuronal function-related pathway

After quality control, 164,684 probes were left for further analysis. We found 4,885 CpG sites were differentially methylated between pretreatment FESs and controls (*t*-test, p < 0.05) as shown in Manhattan plot (Fig.2.A), none of them got the genome-wide significant with p < 1e-6. Most of CpG sites (3,782 out of 4,885, 77.42%) showed hypomethylation in the FESs. We found some top signals with several CpG sites located in one gene, such as the cg12407791 (p=2.03E-05) and cg07010633 (p=1.22E-04) at *UNC13D* (Table 2). The FESs associated CpG sites were enriched in neurogenesis (FDR <1.68e ^−12^), generation of neurons (FDR <1.68e^−12^), and central nervous system development (FDR =4.04e^−10^) (Table 3). Using the replicate dataset, the 637 CpG sites out of the 4,885 were detected, and 6 CpG sites were replicated. They are cg08063724 at the first exon of *MYCL1,* cg17366294 at promoter of *C4orf37*, cg05119831 at gene body of *PRPRN2*, cg00933411 at the promoter of *DLC1,* cg17145652 at 5’UTR of *RNF170* and cg24765079 at body of *CDH1*. We also evaluated the consistency of our findings with previously published SCZ-associated methylation results. Eleven CpG sites found by the largest EWAS in SCZ[7] were replicated in our dataset.

**Figure 2.**
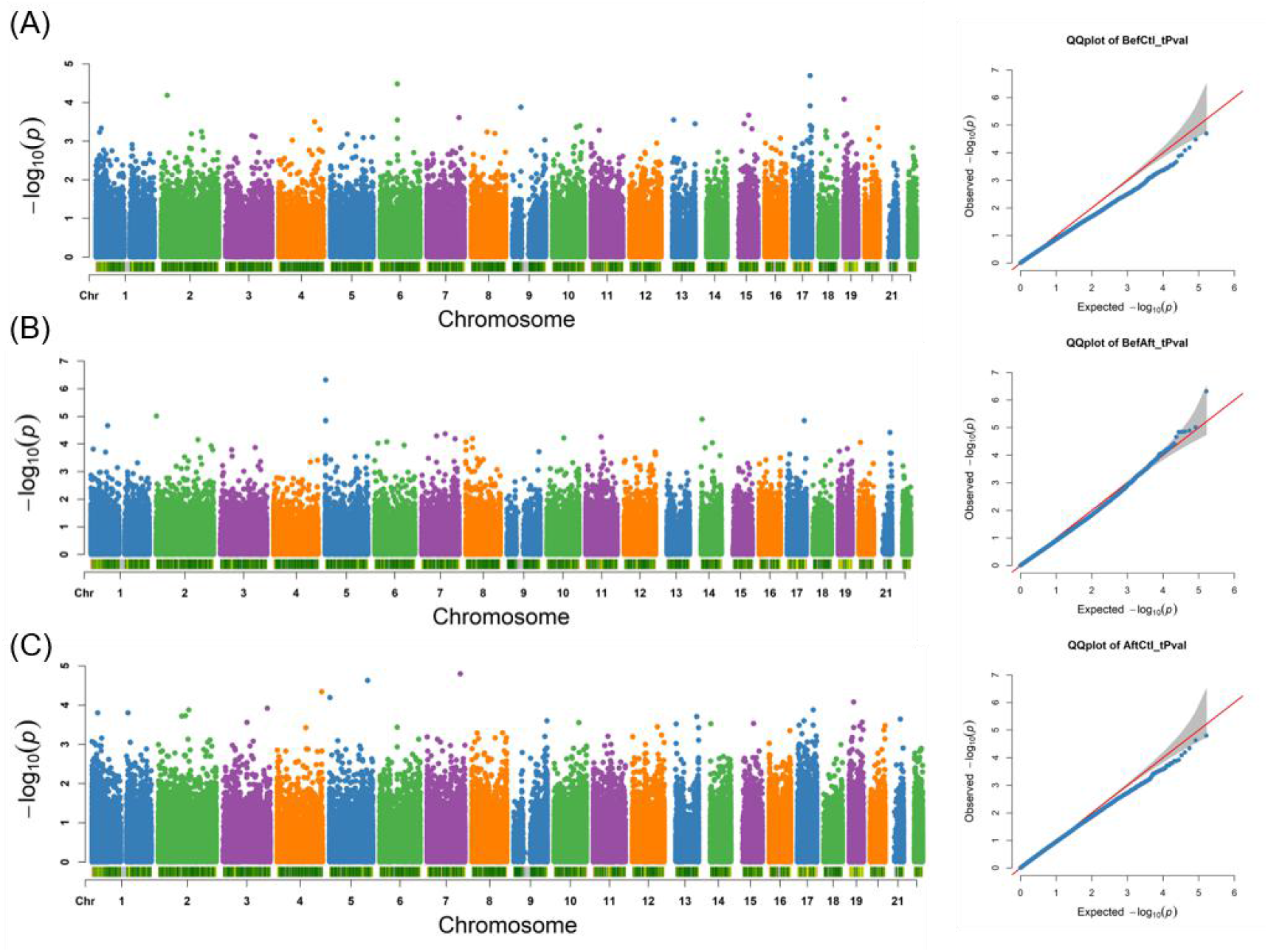
Differential methylation. Manhattan plot and QQ plot of the differentially methylated CpG sites between FES and controls in (A), Pre- and Post-treatment in (B), and post-treatment and control in(C).

We also examined whether the differentially methylated CpG sites are enriched in the 108 loci by GWAS.[28] Among the 108 GWAS loci, there were 64 loci contained 770 CpG sites. In these 770 CpG sites, 77 of them were differentially methylated sites. Enrichment test showed that the GWAS loci had significant enrichment of the differentially methylated CpG sites (Fisher’s Exact test p <1.68e ^−12^). Among the 4,885 differentially methylated CpG sites, 306 CpG sites showed highly correlated methylation between blood and brain using the *Blood Brain DNA Methylation Comparison Tool*.[29]

For the region level analysis, we generated 3,419 regions of gene body and 8,229 regions of promoter from the 164,684 CpG sites tested after quality control. we did not detect pre-treatment FES associated DMRs with P.adj <0.05, but identified 341 gene body DMRs and 858 promoter DMRs with nominal p <0.05.

### 2. Risperidone associated CpG sites and DMRs enriched in calcium signaling pathway

We found one CpG site, cg08778598 at gene body of SDHAP3 genes, was significantly differential methylated between pre-treatment samples and post-treatment samples (P= 4.84e-7) (Fig.2.B). Risperidone treatment increased 3% methylation of this CpG site. There were 5,979 CpG sites with nominal p <0.05 in comparisons of pre-treatment versus post-treatment data, 3,486 (58.61% out of the 5,979 CpG sites) of these CpG sites increased their methylation level after treatment while 2,493 CpG sites decreased the methylation level after treatment. Using replicate data, 517 CpG sites of the 6,142 were detected, and 11 CpG sites were replicated. They are cg09991975 at the promoter of *JOSD1*, cg26309951 at 5’UTR of *MORF4L2*, cg21376883 at body *ACTN2*, cg01598046 at the promoter of *TRAIP*, cg00066816 at the promoter of *IL12B*, cg17617223 at the promoter of *ZER1*, cg15432938 at the promoter of *FRAT1*, cg15875120 at the promoter of *FAM188A*, cg22560190 at the promoter of *CNTN1*, cg12100791 at the promoter of *PYCARD* and cg15928446 at the promoter of *PRR14*.

Pathway analysis showed the 5,979 CpG sites enriched in brain related KEGG pathways such as axon guidance (FDR=1.60E-04), Wnt signaling pathway (FDR = 1.20E-03), MAPK signaling pathway (FDR=1.20E-03) and calcium signaling pathway (FDR=4.84E-03) (Table 3).

For the risperidone-induced differentially methylated regions, we did not detect any DMRs with P.adj < 0.05 in either gene body nor promoter regions, but we detected 210 DMRs in gene body and 555 DMRs in promoter regions with nominal p <0.05.

These genes were functionally enriched for the calcium signaling pathway (FDR=0.0118) and long-term depression (FDR=0.0413). Of the above Entrez genes, the greatest changes were at sites *CACANA1A*, *RYR1*, *NOS1*, *LTB4R2*, and *PTGER3*.

### 3. Post-treatment FES-associated methylation

To help understand the treatment result on the patients, we compared the DNA methylation between post-treatment patients with FES and controls (Fig.2.C). We found 6,760 CpG sites were differentially methylated with nominal p < 0.05. Similar to pre-treatment FES-associated DMPs, most CpG sites (4,034 out of 4,885, 70.00%) showed hypomethylation in post-treatment FES. Using replication data, we detected 3777 of the 6,760 CpG sites and replicated 8 CpG sites. Pathway analysis did not find any KEGG pathways enriched in the 6,760 CpG sites using FDR <0.05, but found several pathways with p <0.05, for example, endocrine resistance (p=1.53e-3), glycosylphosphatidylinositol (GPI)-anchor biosynthesis (p=2.12e-3), insulin signaling pathway (p=4.13e-3), alanine, aspartate and glutamate metabolism (p=2.41e-2), and thyroid hormone signaling pathway(p=2.92e-2).

For the region level analysis, we detected 25 post-treatment FES-associated DMRs in the gene-body with a P.adj < 0.05, including *TP73, PLEKHH3, CNTNAP1, MAD1L1, CHD5, RASA3, GRIN1, VWA5B2, FBXL18, AATK, ZFYVE21, NEURL1B, TRAPPC9, SEMA4C, TBCD, CUX1, LPHN1, CHD3, PRDM16, OBSCN, PLEC1, GALNT9,* and *PTPRN2.* We found 451 DMRs in gene-body regions and 809 DMRs in promoter regions with nominal p < 0.05.

### 4. Risperidone-induced treatment and side effect in methylation

By comparing the direction of effect size between two differential analysis: pre-treatment FEP association analysis (pre-FEP vs control) and treatment association analysis (Post-FEP vs Pre-FEP), we found that 63.02% of the 164,684 CpG sites showed treatment effect with contrasting up-/down-regulation in the two analysis, whereas 36.98% CpG sites showed side effect with common up-/down-regulation in the two analysis (Fig.1.B). For the 4,885 FES-associated CpG sites, 85.32% showed treatment effect, while only 14.68% showed side effect (Fig.3.A). Similar with FES-associated CpG sites, we found the 87.54% out of the Risperidone associated CpG sites showed treatment effect, while only 12.46% showed side effect (Fig.3.B). Different with FES-associated CpG sites and FES-associated CpG sites with majority are treatment effect, Among the 5,979 post-treatment FES associated CpG sites, only 16.84% showed treatment effect, whereas 83.16% showed side effect (Fig.3.C).

**Figure 3.**
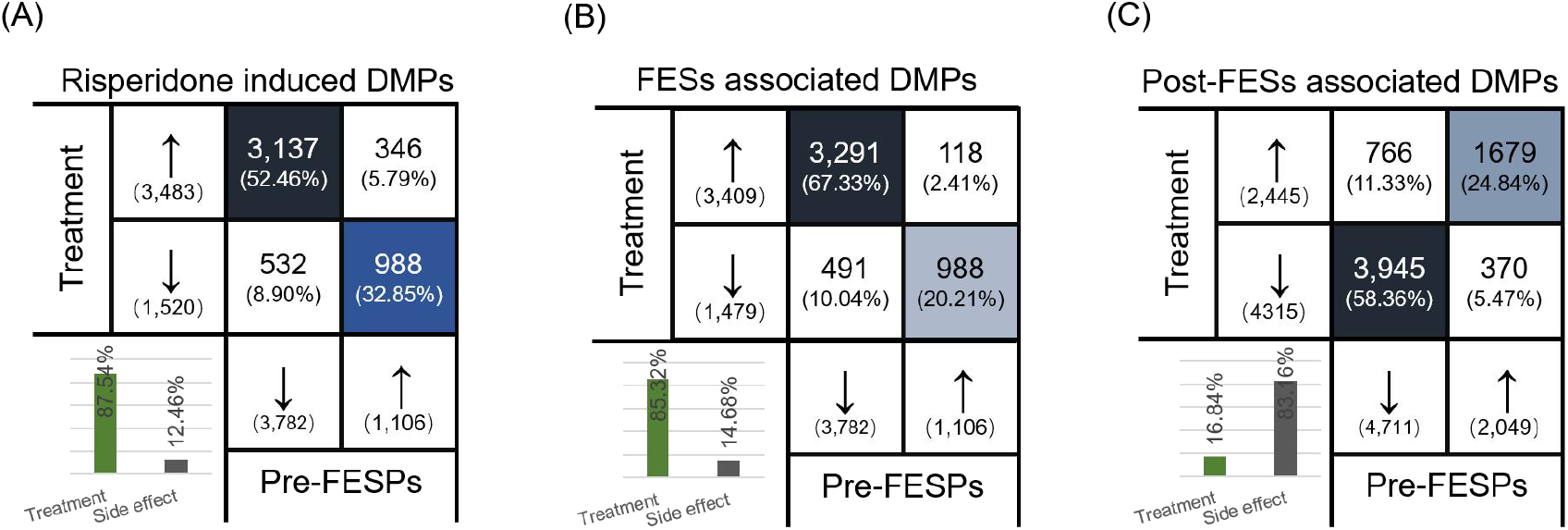
Treatment effect and side-effect. The degree of treatment effect and side effect evident in the CpG sites for FEP-associated CpG sites (A), Risperidone-associated CpG sites (B), and post-treatment FEP associated CpG sites (C).

### 5. Methylation-phenotype network

We further calculated the overlap between the pre-treatment FES associated CpG sites and risperidone associated CpG sites, finding 590 overlapped CpG sites. From these overlapped sites, we also compared post-treatment FES to the healthy controls, identifying 580 normalized CpG sites. These 590 CpG sites mapped to 568 unique genes which contained 113 SCZ related candidate genes obtained from SZDB (http://www.szdb.org/) and *NPdenovo* (http://www.wzgenomics.cn/NPdenovo/), such as MAN2A1 (Supplementary Table S2).

To explore whether these normalized CpG sites can reflect the phenotypic changes in the brain, we correlated methylation changes to the phenotypic changes. They ranged from symptom severity, to cognitive function, to brain structures. We then found 284 out of the 580 normalized CpG sites that correlated with at least one phenotype (absolute correlation coefficient > 0.3, p <0.05, Supplementary Figure S1). These included two CpG sites (cg09175724 and cg19248041) correlated with five phenotypic variables, nine CpG sites correlated with four phenotypic variables, 25 CpG sites correlated with three phenotypic variables, and 61 CpG sites correlated with two phenotypic variables. For example, cg09175724 at the 5’UTR of CDC42EP2, correlated with reducing rate of PANSS-Total score (correlation coefficient =0.40, p= 1.30E-02), changes of PANSS-Positive symptom score (correlation coefficient =0.57, p= 2. 07E-04), changes of PANSS-General symptom score (correlation coefficient =0.36, p= 2. 50E-02), and reducing rate of fALFF (resting state fMRI) in both left and right putamen (correlation coefficient =−0. 39, p= 1.30E-02).

For the phenotypic variables, we found methylation changes after treatment in 96 sites correlating with changes in PANSS scores (Figure S2); in 129 sites, methylation changes correlated with reduced cognitive function; and in 122 sites, methylation changes were correlated with changes in spontaneous brain activity found in fMRI (Supplementary Figure S3) (absolute Spearman correlation coefficient > 0.3, p < 0.05).

Methylation of one CpG site (cg25535999) at the gene body of NR3C1, which encodes the glucocorticoid receptor, was normalized by treatment, and changes in that methylation correlated with PANSS-G, PANSS-T changes and cognitive improvement measured by the SCWT. Another CpG site, cg25114611 at the promoter of FKBP5 (TSS1500) was also normalized after treatment. Methylation changes of cg25114611 correlated with changes of nodal degree of anterior cingulate and paracingulate gyri (ACG, left and right) as well as with changes of nodal efficiency of right ACG. We found 11 normalized CpG sites in calcium pathway genes correlated with multiple phenotypic changes. For example, cg06204009 at the promoter of ATP2B3 and cg17119907 at the 5’UTR of NOS1 were normalized after treatment and correlated with the PANSS-G changes. In the calcium pathway, CpG site cg26571093 at the gene body of CACNA1H was normalized and correlated with changes in resting state brain function within the caudate nucleus (left and right) and with cognitive function improvement measured by the SCWT (Supplementary Table S3).

## Discussion and conclusion

This is the first study of treatment-naïve patients with FES now receiving monotherapy with risperidone to examine methylation changes in peripheral blood to reflect treatment and side effect. This is also the first study to examine such changes as they relate to neuroimaging and cognitive phenotypes within this population. Through comprehensive comparisons between pre- and post-treatment FES populations and controls samples, we found treatment effect and side effect at the molecular level and identified hundreds of normalized CpG sites that parallel brain phenotypic changes.

Limiting our study to the FES population avoided the potentially confounding effects of chronic illness and drug treatment. This is important for both of our aims: 1) identifying disease-related methylation changes before treatment, and 2) investigating acute antipsychotic treatment effects on DNA methylation. Although our sample size is small due to the limited population, our findings in regard to these two aims are well supported by published research. The over-representation of FES-associated CpG sites within the SCZ-related GWAS loci is consistent with the largest SCZ EWAS study[7]. In regard to the treatment effect on DNA methylation, some of the associated genes are pivotal to neuronal function (e.g., *HDAC6*), while some have been previously associated with SCZ by GWASs (e.g., *MAN2A1, CNTN4, MEF2C*), and others are involved in *de novo* mutations related to SCZ (e.g., *NCKAP1, CAPRIN1, DNMT3A*). Several of the genes are also implicated in long-term synaptic depression and disease-associated calcium signaling pathways. Several evidences have implicated dysfunctional of these pathways in the pathophysiology of SCZ[30].

Notably, the use of monotherapy, consistent drug exposure period, and consistent dosage enhanced the validity of the pre- and post-treatment comparisons. Furthermore, having healthy controls as a reference allowed us to distinguish treatment effect from side effects by equating changes in DNA methylation with common up/down regulation. We found majority of the risperidone induced DNA methylation changes had treatment effect. This result supports the clinal studies that antipsychotic drugs do more good than harm[31] for the patients in a molecular aspect. The comparison between the pre-treatment FES associated CpG sites and the post-treatment associated CpG sites showed us different effect pattern with majority of treatment effect in the pre-treatment FES and majority of side effect in the post-treatment FES. It means that the differences between the treated patients verse controls are more likely the drug induced difference instead of the disease induced methylation changes. These findings suggested that when we try to identify the new drug target using the drug naïve patients instead of the treated patients is critical to find the real signals.

For the first time in schizophrenia research, we linked post-antipsychotic methylation changes with changes in psychiatric symptoms as well as neuropsychological and neurophysiological abnormalities in patients with schizophrenia. We relate the methylation-phenotype network identified in this study to brain changes for two reasons. First, methylation status and methylation changes induced by the drug in blood cells correlate closely with methylation in the brain.[16, 32]. Second, while previous studies in schizophrenia associated methylation with changes in cognitive effect and MRI[33], we linked post-antipsychotic methylation changes with changes in psychiatric symptoms. The overlap of DNA methylation and established schizophrenia risk loci is of significant clinical relevance. Studies of methylation in blood cells may provide an important approach for understanding SCZ pathophysiology and drug treatment effectiveness.

Certain limitations of our clinical study are worth noting. Although our conclusions are supported statistically and were to a degree validated in an independent dataset, the results need to be validated in a larger cohort with FES. Future studies could identify the time course of methylation changes after therapy with several important clinical implications. Rapid onset of methylation may be a useful index of future treatment response and therefore guide earlier changes in drug therapy for treatment non-responders. Variability in timing or extent of methylation changes across patients might account for variability in the speed and extent of treatment effect. Methylation changes that diminish or persist after longer term treatment might be related to course of illness, risk for relapse, etc. Last but not least, we measured the treatment and side effect in the peripheral blood which may not directly reflect the effect in the brain. Cause the brain tissue is impossible to get for these longitudinal studies, further animal model and cell lines studies are need.

## Materials and Methods

### Participants

Participants included 38 right-handed Chinese Han patients with FES (25 men; 13 women; mean age, 25.0 years; age range, 18-37) and 38 demographically-matched healthy controls (25 men; 13 women; mean age, 24.8 years; age range 18-32) (Table 1). Two attending psychiatrists diagnosed participants using the Structured Clinical Interview for DSM-IV-TR, patient version (SCID-I/P). All patients with FES had an onset of psychotic symptoms less than one-year before study participation. Healthy controls, had no history of Axis I or II disorders (Structured Clinical Interview for DSM-IV Axis I Disorders—Non-Patient Edition (SCID-I/NP); Structured Clinical Interview for DSM-IV Axis II Disorders), nor any known first-degree family history of psychiatric illness. Individuals with neurological disease, systemic disease or substance abuse disorders (such as Alcohol use disorders[34]) were excluded from this study.

**Table 1.**
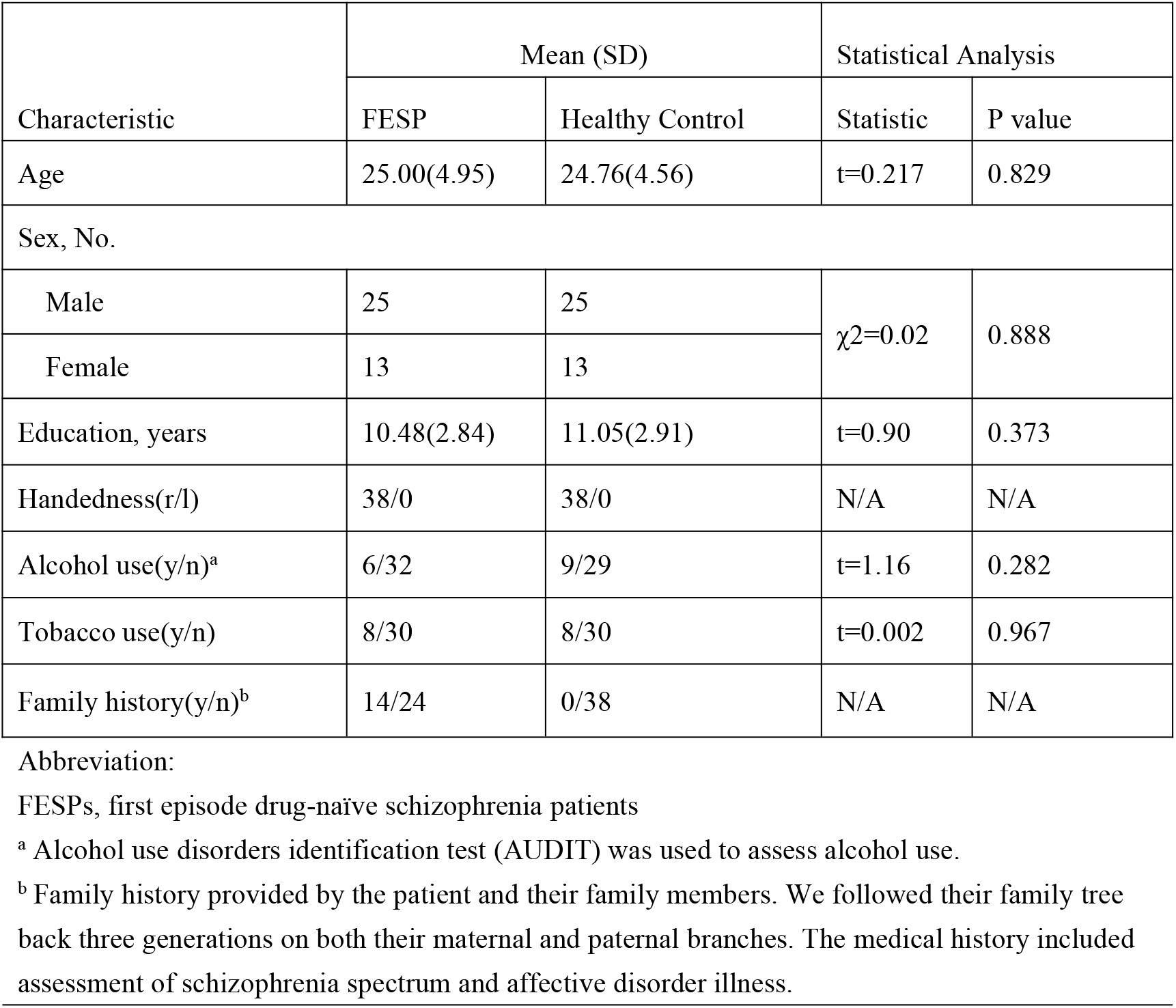
Demographic and Clinical Characteristics of Drug-Naïve First Episode Schizophrenia Patients and Matched Healthy Control Participants

**Table 2.** Top signals for FESPs associated differential methylation and risperidone-induced methylation

| Probe information |  |  |  |  | Statistical Analysis |  |  |
| --- | --- | --- | --- | --- | --- | --- | --- |
| Probe | Chr | Position | Gene<sup>a</sup> | Feature<sup>b</sup> | t/ paired t<sup>c</sup> | P value | $\Delta^d$ |
| FESPs associated differential methylation |  |  |  |  |  |  |  |
| cg12407791 | 17 | 73824354 | UNC13D | Body | -4.556 | 2.03E-05 | -0.046 |
| cg12070285 | 6 | 74072447 | C6orf221 | 1stExon | -4.469 | 3.32E-05 | -0.014 |
| cg00401265 | X | 78003113 | LPAR4 | Promoter | 4.349 | 4.31E-05 | 0.024 |
| cg13918808 | X | 13062651 | FAM9C | 5'UTR | 4.272 | 6.03E-05 | 0.018 |
| cg15150970 | 2 | 25473529 | DNMT3A | Body | -4.247 | 6.57E-05 | -0.029 |
| cg15226147 | 19 | 1275266 | C19orf24 | Promoter | -4.169 | 8.23E-05 | -0.013 |
| cg25289658 | X | 135278646 | FHL1 | 5'UTR | -4.105 | 1.03E-04 | -0.048 |
| cg03598919 | X | 51150887 | CXorf67 | 1stExon | 4.101 | 1.12E-04 | 0.017 |
| cg07010633 | 17 | 73824396 | UNC13D | Body | -4.067 | 1.22E-04 | -0.063 |
| cg13728834 | 9 | 35846324 | TMEM8B | Body | -4.035 | 1.32E-04 | -0.035 |

| Risperidone-induced methylation changes |  |  |  |  |  |  |  |
| --- | --- | --- | --- | --- | --- | --- | --- |
| cg08778598 | 5 | 1594579 | SDHAP3 | Body | -6.084 | 4.84E-07 | 0.028 |
| cg04174838 | X | 3733239 | LOC389906 | IGR | -5.265 | 6.21E-06 | 0.027 |
| cg01445411 | 2 | 1358684 | SNTG2 | Body | -5.117 | 9.83E-06 | 0.013 |
| cg25396787 | X | 3733991 | LOC389906 | IGR | -5.106 | 1.02E-05 | 0.031 |
| cg10193721 | 14 | 24780691 | LTB4R2 | Body | -5.029 | 1.29E-05 | 0.026 |
| cg06721546 | 5 | 1594808 | SDHAP3 | Promoter | -5.003 | 1.40E-05 | 0.025 |
| cg05571310 | 17 | 72350354 | KIF19 | Body | -4.995 | 1.43E-05 | 0.016 |
| cg27149073 | 5 | 1594330 | SDHAP3 | Body | -4.988 | 1.46E-05 | 0.038 |
| cg23823908 | 1 | 71512485 | PTGER3 | 1stExon | -4.861 | 2.17E-05 | 0.026 |
| cg19761502 | 21 | 40759534 | WRB | Body | -4.673 | 3.86E-05 | 0.018 |
| cg11633514 | 7 | 100183910 | LRCH4 | Promoter | -4.634 | 4.35E-05 | 0.006 |
| cg15949805 | 7 | 63505714 | ZNF727 | Promoter | -4.579 | 5.14E-05 | 0.019 |
| cg08234507 | 11 | 64990959 | SLC22A20 | Body | 4.554 | 5.55E-05 | -0.010 |
| cg17477052 | 10 | 71596731 | COL13A1 | Body | -4.525 | 6.06E-05 | 0.035 |
| cg03066788 | 8 | 28175285 | PNOC | 5'UTR | -4.501 | 6.50E-05 | 0.032 |
| cg14771419 | 7 | 142012988 | PRSS58 | IGR | -4.500 | 6.54E-05 | 0.028 |
| cg01047586 | 2 | 176933407 | EVX2 | IGR | -4.471 | 7.13E-05 | 0.020 |
| cg23958373 | 8 | 599963 | ERICH1 | IGR | 4.414 | 8.48E-05 | -0.025 |
| cg22248011 | 6 | 52442381 | TRAM2 | Promoter | -4.414 | 8.48E-05 | 0.005 |
| cg07234876 | 8 | 600039 | ERICH1 | IGR | 4.409 | 8.59E-05 | -0.031 |
| cg10616337 | 20 | 5986410 | CRLS1 | Promoter | 4.405 | 8.70E-05 | -0.012 |
| cg18306861 | 14 | 67879101 | PLEK2 | Promoter | -4.389 | 9.13E-05 | 0.017 |

Abbreviations: Chr, chromosome; FESPs, first episode drug naïve schizophrenia patients
<sup>a</sup> Gene name with Illumina gene annotation
<sup>b</sup> Indicates if the loci overlap with untranslated region(UTR), first Exon (1stExon),gene body, intergenic region (IGR) or promoter (upstream of transcription start site,)
<sup>c</sup> For FESPs associated differential methylation, it is t value, for risperidone-induced differential methylation, it is paired t value.
<sup>d</sup> The mean difference between patients and controls, for FESPs associated differential methylation, is the mean methylation value of FESPs minus CTLs. For risperidone-induced methylation change, it is the mean methylation value of post-treatment minus pre-treatment

**Table 3.** Functional annotation for FESPs associated differential methylation, Risperidone-induced differential methylation and normalized methylation genes

| Gene set | Description | C<sup>a</sup> | O<sup>b</sup> | P Value | FDR corrected P Value |
| --- | --- | --- | --- | --- | --- |
| FESPs associated differential methylation |  |  |  |  |  |
| GO |  |  |  |  |  |
| GO:0022008 | neurogenesis | 1414 | 216 | 0.00E+00 | 0.00E+00 |
| GO:0048699 | generation of neurons | 1319 | 204 | 0.00E+00 | 0.00E+00 |
| GO:0030182 | neuron differentiation | 1197 | 188 | 3.33E-16 | 1.68E-12 |
| GO:0007417 | central nervous system development | 890 | 144 | 1.33E-13 | 4.04E-10 |
| GO:2000026 | regulation of multicellular organismal development | 1675 | 225 | 9.14E-12 | 2.30E-08 |
| Risperidone-induced differential methylation |  |  |  |  |  |
| KEGG |  |  |  |  |  |
| hsa04360 | Axon guidance | 177 | 66 | 5.30E-07 | 1.60E-04 |
| hsa04310 | Wnt signaling pathway | 143 | 53 | 8.51E-06 | 1.20E-03 |
| hsa04010 | MAPK signaling pathway | 255 | 83 | 1.19E-05 | 1.20E-03 |
| hsa04390 | Hippo signaling pathway | 154 | 54 | 4.32E-05 | 2.62E-03 |
| hsa04550 | Signaling pathways regulating pluripotency of stem cells | 142 | 50 | 7.27E-05 | 3.67E-03 |
| hsa04020 | Calcium signaling pathway | 182 | 60 | 1.28E-04 | 4.84E-03 |
| hsa04144 | Endocytosis | 260 | 77 | 7.24E-04 | 1.99E-02 |
| GO |  |  |  |  |  |
| GO:0007409 | axonogenesis | 417 | 160 | 0.00E+00 | 0.00E+00 |
| GO:0007417 | central nervous system development | 890 | 327 | 0.00E+00 | 0.00E+00 |
| GO:0007420 | brain development | 668 | 240 | 0.00E+00 | 0.00E+00 |
| GO:0009790 | embryo development | 945 | 328 | 0.00E+00 | 0.00E+00 |
| GO:0022008 | neurogenesis | 1414 | 490 | 0.00E+00 | 0.00E+00 |
| GO:0030182 | neuron differentiation | 1197 | 422 | 0.00E+00 | 0.00E+00 |
| GO:0030900 | forebrain development | 358 | 157 | 0.00E+00 | 0.00E+00 |
| Normalized methylation genes |  |  |  |  |  |
| GO |  |  |  |  |  |
| GO:0022008 | neurogenesis | 1414 | 123 | 1.84E-14 | 1.58E-10 |
| GO:0048699 | generation of neurons | 1319 | 116 | 5.62E-14 | 2.40E-10 |
| GO:0030182 | neuron differentiation | 1197 | 107 | 2.25E-13 | 4.81E-10 |
| GO:0048666 | neuron development | 951 | 86 | 3.87E-11 | 6.61E-08 |
| GO:0031175 | neuron projection development | 811 | 73 | 1.65E-09 | 2.10E-06 |
| GO:0048667 | cell morphogenesis involved in neuron differentiation | 506 | 51 | 1.50E-08 | 1.21E-05 |
| GO:0048812 | neuron projection morphogenesis | 552 | 54 | 1.56E-08 | 1.21E-05 |
| GO:0051960 | regulation of nervous system development | 760 | 65 | 9.54E-08 | 5.44E-05 |
| GO:0007417 | central nervous system development | 890 | 72 | 1.70E-07 | 8.55E-05 |
| GO:0007267 | cell-cell signaling | 1557 | 109 | 2.07E-07 | 9.83E-05 |

Abbreviations:
<sup>a</sup> Total number of genes in the term.
<sup>b</sup> Number of genes in the term with overlap in FESPs associated differential methylation,
Risperidone-induced differential methylation and normalized gene methylation

All procedures were approved by the ethics committee of the Second Xiangya Hospital and the Second Affiliated Hospital of Xinxiang Medical University. All participants provided written informed consent and could discontinue study participation at any time.

### Medication, psychiatric assessments and neuroimaging phenotyping

The SCZ cohort was treated with risperidone monotherapy at a dosage of 4mg to 6 mg/day for eight weeks, without the addition of mood stabilizers or antidepressants. Symptom severity was evaluated at baseline and follow-up with the Positive and Negative Syndrome Scale (PANSS)[35] scored to generate total score (PANSS-T), positive symptom score (PANSS-P), negative symptom score (PANSS-N) and the PANSS general psychopathological symptom score (PANSS-G). Three of them, PANSS-T, PANSS-P and PANSS-G, significantly improved with treatment (paired t test p < 0.001).

Magnetic resonance imaging (MRI) scans were performed on all participants using a 3T MRI scanner (Siemens Healthcare; Erlangen, Germany) with a 16-channel head coil. Diffusion tensor imaging, T1-weighted imaging and resting-state fMRI data were collected at baseline and again after 8 weeks. Spontaneous brain activity was quantified using the fractional amplitude of low-frequency fluctuations (fALFF) and regional homogeneity (ReHo). Detailed scanning parameters and analysis method for these MRI data were performed as previously described.[36, 37] Cognitive function was evaluated in parallel with MRI scanning using standardized and widely-used neuropsychological tests: Stroop Color Word Test (SCWT), Wisconsin Card Sorting Test (WCST), Trail Making Test (TMT), Verbal Fluency Test (VFT), and Digit Span Distraction Test (DSDT).[38–40]

All phenotypic variables (MRI, Cognition) used in this analysis were from brain regions or cognitive measures that changed significantly between pre- and post-treatment in the SCZ cohort or were significantly different between cohorts (supplementary Table S1A-S1C). Significantly improved phenotypes (paired t test p < 0.001) were included in the analyses to determine the relation between methylation changes and phenotypic improvement; changes observed after treatment were generally consistent with earlier literature.[41–47] For spontaneous brain activity, we included fALFF in bilateral putamen and the right caudate and ReHo in the right caudate and left putamen. For cognitive function, we included changes of SCWT, WSCT, and TMT. Statistical analysis examining these phenotypic changes in relation to methylation changes are shown in supplementary Table S1A, S1B, and S1C.

### Quantification and analysis of DNA methylation

#### Microarray processing

We collected blood samples from 38 healthy controls at baseline and 38 patients at two time-points (i.e., before and after risperidone treatment). Whole genome methylation status was then examined in 114 samples. DNA (500ng) was isolated using QIAamp DNA Blood Mini Kit (Qiagen; Germantown, MD) and treated with sodium bisulfite using the EZ DNA methylation kit (Zymo Research; Irvine, CA). DNA methylation was quantified using Infinium^®^ Human Methylation 450K BeadChip (Illumina Inc.; San Diego, CA).

#### Data processing

All analyses were performed in R version 3.3.1. Raw intensity files were preprocessed and quantiles were normalized using the Bioconductor package ChAMP, version 2.0.1.[48] Proportions of methylation values (Illumina “Beta” scale) were calculated. After which, BMIQ[49] were used to adjust for type II bias. Probes were removed according to the following criteria: 1) detection *p*-value above 0.01 in one or more samples; 2) bead counts less than 3 in at least 5% of samples; 3) having SNPs; 4) aligning to multiple locations; and 5) identified in Nordlund *et al.*[50]; and 6) located in sex chromosomes. There were 164,684 probes remaining for differential methylation analysis. Batch and positional effect of each chip was adjusted using the ComBat empirical Bayesian approach *[51]*. The reference-based method was used to calculate cell type proportions[52]. We used linear regression to regress out effects of cell type proportion, age, sex, smoking status, and drinking status for each probe.

#### Power analysis

Given a sample size of 76 for the risperidone-association analysis, there was 80% power to detect 12% mean methylation changes at P < 1e-6. For the case control analysis of 76 samples, there was 80% power to detect 13% mean methylation changes at P < 1e-6. The power analysis was based on the power simulations across a range of sample sizes and effect sizes according to the calculation of Saffari et al[53].

#### Identification of differentially methylated individual CpG sites and genomic regions

To identify differentially methylated individual CpG sites, we conducted three epigenome-wide association analysis: the FES association analysis on pre-treatment FES verse controls, the risperidone association analysis on pre-treatment FES verse post-treatment FES, and the post-treatment FES association analysis on post-treatment FES verse control (Fig.1). Linear regression model in *Limma* package[54] was used for testing the differential DNA methylation between pre-/post-treatment FES and controls at each CpG site. Paired *t*-tests were used to test for differential DNA methylation between pre- and post-treatment samples in the patients. For multiple testing correction, we followed the Bonferroni correction and used the threshold of 1e-6 as recommended by Saffari *et al.* [53] and Rakyan *et al.[8]*. We reported the results with nominal (p=0.05) and genome-wide (p=1e-6) significance levels.

To identify differentially methylated regions (DMRs), we used the mCSEA (Methylated CpGs Set Enrichment Analysis)[55] in R (version 3.5). We ran the mCSEA analysis for pre-treatment FES versus post-treatment FES (paired test), pre-treatment FES versus control samples, and post-treatment FES versus control samples. We ranked all evaluated CpG sites after quality control with the differential statistics above. Then we used the *mCSEATest* function to search promoter and gene body DMRs. We specified five CpGs as the minimum amount per region and performed 10000 permutations to calculate P value as the default. The p value was then adjusted by the Benjamini and Hochberg procedure for multiple testing correction.

#### Estimation of risperidone-induced treatment and side effect in DNA methylation

To estimate treatment and side effect induced by risperidone in DNA methylation, we compared the direction of effect size between two differential analysis: pre-treatment FEP association analysis (pre-FEP vs control) and treatment association analysis (Post-FEP vs Pre-FEP) (Fig.1.B). In pre-treatment FEP association analysis, we defined up-regulated CpG sites as CpG sites that have higher methylation levels in patients than in controls, while the down-regulated CpG sites we defined as lower methylation levels in patients than in controls. In the treatment association analysis, the up-regulated CpG sites were defined as CpG sites with higher methylation levels in post-treatment patients then pre-treatment patients, while the down-regulated CpG sites had lower methylation levels in post-treatment patients than in pre-treatment patients.

Based on the two comparisons we divided the CpG sites into two groups: the CpG sites with common up/down regulation, and CpG sites with contrasting up/down regulation. Any changes in methylation in the contrasting CpG sites represent treatment effect. For example, before treatment a CpG site is down regulated in patients but not in controls; whereas after treatment, the methylation levels rise making them closer to normal. In the same way, the changes in CpG sites with common up/down regulation represent side effect, meaning that with side-effect the treatment makes the methylation levels in the patients even farther from the normal control.

To characterize the degree of treatment effect and side effect evident in the CpG sites, we estimated the percentages of CpG sites with common up/down regulation and contrasting up/down regulation for each of the following categories:

1. Total CpG sites after quality control, representing the overall effect of treatment on DNA methylation
2. FEP-associated CpG sites—dysfunctional CpG sites or genes
3. Risperidone-associated CpG sites
4. Post-treatment FEP associated CpG sites—final results of treatment.

#### Over-representation analysis

To evaluate the enrichment of differential DNA methylation in SCZ GWAS signals, we downloaded the 108 loci from PGC website[28]. For every GWAS region, we counted the number of probes detected in this study and the number of the FES associated differentially methylated probes. SCZ related candidate genes were also obtained from SZDB (http://www.szdb.org/)[56] and *NPdenovo* (http://www.wzgenomics.cn/NPdenovo/)*[57]*. To evaluate the enrichment of differential DNA methylation with CpG sites that have brain-blood correlated methylation, we downloaded the CpG sites list from *Blood Brain DNA Methylation Comparison Tool* (http://epigenetics.essex.ac.uk/bloodbrain/). A two-sided Fisher’s exact test was used to estimate the enrichment. Gene ontology and KEGG pathway analyses were performed using *missMethyl* package, which consider the number of CpG islands corresponding to each gene[58].

#### Replication

We obtained a replication data set involving participants from a prior study involving similar entry criteria (untreated first episode schizophrenia), treatment (4-6 weeks risperidone), and phenotyping[59]. This sample included four controls (2 men, 2 women; 2 African American, 1 Hispanic and 1 White; age range, 27-30) and three FESs (1 men, 2 women; 2 African American, 1 Hispanic; age range, 18-40) (Supplementary Table S1D). For each subject, blood samples were isolated twice pre- and post-treatment (six weeks). DNA methylation was quantified using Infinium Human Methylation 27 BeadChip (Illumina, CA) at the Northwestern University Core facility.

### Construction of the methylation-phenotype network

To estimate parallel DNA methylation changes and phenotype, we constructed a methylation-phenotype network by correlating the changes of DNA methylation and phenotypic variables. We choose the CpG sites with normalized effect to construct the network analysis. The normalized CpG sites were those overlapped between FES-associated CpG sites and treatment induced CpG sites, excluded the post-FESs associated CpG sites. All the phenotypic variables used were listed in the Supplementary Table S1 based on previous publications.[36, 37] The relationships between pairs of variables (for example, methylation change of a CpG site and change of PANSS scores) were examined using Spearman rank correlation tests. Associations with an absolute correlation coefficient larger than 0.3 and a *p*-value smaller than 0.05 were plotted in the network, and these variables were treated as edges in the graph (Figure 2). The graph was made using Cytoscape[60] v3.5.0 (http://cytoscape.org/) software. False discovery rate (FDR) was used to correct for multiple comparisons. Correlations with a FDR corrected p < 0.05 were considered significant.

## Acknowledgments

This work was supported by grants from the National Natural Science Foundation of China (81871057 and 81371480 to J.T;81271484 and 81471361 to X.C.; 31970572, 81401114 and 31571312 to C.C.) the National Key Plan for Scientific Research and Development of China (2016YFC1306000) (to C. C.) and NIH grant 1 U01 MH103340-01, 1R01ES024988 (to C. L.), MH083888 (to J.R.B.). Wuhan Science and Technology Bureau grant (2017060201010169 to M. H.). Central South University Graduate Project grant (502221702 to Y.X.).

Dr Hu, Dr Zong and Ms Xia had full access to all the data. Ms Xia takes responsibility for the integrity of the data and the accuracy of the data analysis. Dr. J. Tang, Dr X. Chen, Dr Liu and Dr C. Chen supervised this work and contributed equally as principal investigators.

## Study concept and design

Jinsong Tang, Xiaogang Chen, Chunyu Liu, Chao Chen

## Acquisition data

Maolin Hu, Xiaofen Zong, Jinsong Tang, Jeffrey R Bishop, Zongchang Li, Ying He, Yanhui Liao.

## Analysis and interpretation of data

Yan Xia, Chao Chen, John Sweeney, Chunyu Liu, Leah Rubin, Bingshan Li, Jinsong Tang, Yanhui Liao, Xiaogang Chen.

## Drafting of the manuscript

Yan Xia.

## Critical revision of the manuscript for important intellectual content

Yan Xia, John Sweeney, Jinsong Tang, Chunyu Liu, Chao Chen, Yanhui Liao, Yunpeng Wang, Gina Giase, Jeffrey Bishop, Liz Kunney.

## Statistical analysis

Yan Xia, Chao Chen, Bingsha Li, Jinsong Tang

## Obtained funding

Jinsong Tang, Xiaogang Chen, Chao Chen, Chunyu Liu, Maolin Hu, Yan Xia, Jeffrey Bishop.

## Study supervision

Jinsong Tang, Xiaogang Chen, Chao Chen, Chunyu Liu.

Dr. Sweeney consulted to Takeda. None of the other authors report potential conflicts of interest.

## Reference

1. Miyamoto S, Duncan GE, Marx CE, Lieberman JA: Treatments for schizophrenia: a critical review of pharmacology and mechanisms of action of antipsychotic drugs. Mol Psychiatry 2005, 10(1):79–104.

2. Tiihonen J, Mittendorfer-Rutz E, Majak M, Mehtala J, Hoti F, Jedenius E, Enkusson D, Leval A, Sermon J, Tanskanen A et al: Real-World Effectiveness of Antipsychotic Treatments in a Nationwide Cohort of 29823 Patients With Schizophrenia. JAMA Psychiatry 2017.

3. Thomas EA: Molecular profiling of antipsychotic drug function: convergent mechanisms in the pathology and treatment of psychiatric disorders. Mol Neurobiol 2006, 34(2):109–128.

4. Thomas EA, George RC, Danielson PE, Nelson PA, Warren AJ, Lo D, Sutcliffe JG: Antipsychotic drug treatment alters expression of mRNAs encoding lipid metabolism-related proteins. Mol Psychiatry 2003, 8(12):983–993, 950.

5. Langlois MC, Beaudry G, Zekki H, Rouillard C, Levesque D: Impact of antipsychotic drug administration on the expression of nuclear receptors in the neocortex and striatum of the rat brain. Neuroscience 2001, 106(1):117–128.

6. Chong VZ, Young LT, Mishra RK: cDNA array reveals differential gene expression following chronic neuroleptic administration: implications of synapsin II in haloperidol treatment. J Neurochem 2002, 82(6):1533–1539.

7. Hannon E, Dempster E, Viana J, Burrage J, Smith AR, Macdonald R, St Clair D, Mustard C, Breen G, Therman S et al: An integrated genetic-epigenetic analysis of schizophrenia: evidence for co-localization of genetic associations and differential DNA methylation. Genome biology 2016, 17(1):176.

8. Rakyan VK, Down TA, Balding DJ, Beck S: Epigenome-wide association studies for common human diseases. Nat Rev Genet 2011, 12(8):529–541.

9. Jaffe AE, Gao Y, Deep-Soboslay A, Tao R, Hyde TM, Weinberger DR, Kleinman JE: Mapping DNA methylation across development, genotype and schizophrenia in the human frontal cortex. Nat Neurosci 2016, 19(1):40–47.

10. Liu J, Chen J, Ehrlich S, Walton E, White T, Perrone-Bizzozero N, Bustillo J, Turner JA, Calhoun VD: Methylation patterns in whole blood correlate with symptoms in schizophrenia patients. Schizophr Bull 2014, 40(4):769–776.

11. Pidsley R, Viana J, Hannon E, Spiers H, Troakes C, Al-Saraj S, Mechawar N, Turecki G, Schalkwyk LC, Bray NJ et al: Methylomic profiling of human brain tissue supports a neurodevelopmental origin for schizophrenia. Genome Biol 2014, 15(10):483.

12. Kinoshita M, Numata S, Tajima A, Shimodera S, Ono S, Imamura A, Iga J, Watanabe S, Kikuchi K, Kubo H et al: DNA methylation signatures of peripheral leukocytes in schizophrenia. Neuromolecular Med 2013, 15(1):95–101.

13. Nishioka M, Bundo M, Koike S, Takizawa R, Kakiuchi C, Araki T, Kasai K, Iwamoto K: Comprehensive DNA methylation analysis of peripheral blood cells derived from patients with first-episode schizophrenia. J Hum Genet 2013, 58(2):91–97.

14. Dempster EL, Pidsley R, Schalkwyk LC, Owens S, Georgiades A, Kane F, Kalidindi S, Picchioni M, Kravariti E, Toulopoulou T et al: Disease-associated epigenetic changes in monozygotic twins discordant for schizophrenia and bipolar disorder. Hum Mol Genet 2011, 20(24):4786–4796.

15. Mill J, Tang T, Kaminsky Z, Khare T, Yazdanpanah S, Bouchard L, Jia P, Assadzadeh A, Flanagan J, Schumacher A et al: Epigenomic profiling reveals DNA-methylation changes associated with major psychosis. Am J Hum Genet 2008, 82(3):696–711.

16. Aberg KA, McClay JL, Nerella S, Clark S, Kumar G, Chen W, Khachane AN, Xie L, Hudson A, Gao G et al: Methylome-wide association study of schizophrenia: identifying blood biomarker signatures of environmental insults. JAMA Psychiatry 2014, 71(3):255–264.

17. Montano C, Taub MA, Jaffe A, Briem E, Feinberg JI, Trygvadottir R, Idrizi A, Runarsson A, Berndsen B, Gur RC et al: Association of DNA Methylation Differences With Schizophrenia in an Epigenome-Wide Association Study. JAMA Psychiatry 2016, 73(5):506–514.

18. Tang H, Dalton CF, Srisawat U, Zhang ZJ, Reynolds GP: Methylation at a transcription factor-binding site on the 5-HT1A receptor gene correlates with negative symptom treatment response in first episode schizophrenia. Int J Neuropsychopharmacol 2014, 17(4):645–649.

19. Guidotti A, Grayson DR: DNA methylation and demethylation as targets for antipsychotic therapy. Dialogues Clin Neurosci 2014, 16(3):419–429.

20. Melka MG, Castellani CA, Laufer BI, Rajakumar RN, O’Reilly R, Singh SM: Olanzapine induced DNA methylation changes support the dopamine hypothesis of psychosis. J Mol Psychiatry 2013, 1(1):19.

21. Melka MG, Laufer BI, McDonald P, Castellani CA, Rajakumar N, O’Reilly R, Singh SM: The effects of olanzapine on genome-wide DNA methylation in the hippocampus and cerebellum. Clin Epigenetics 2014, 6(1):1.

22. Melka MG, Castellani CA, Rajakumar N, O’Reilly R, Singh SM: Olanzapine-induced methylation alters cadherin gene families and associated pathways implicated in psychosis. BMC Neurosci 2014, 15:112.

23. Murata Y, Nishioka M, Bundo M, Sunaga F, Kasai K, Iwamoto K: Comprehensive DNA methylation analysis of human neuroblastoma cells treated with blonanserin. Neurosci Lett 2014, 563:123–128.

24. Melka MG, Rajakumar N, O’Reilly R, Singh SM: Olanzapine-induced DNA methylation in the hippocampus and cerebellum in genes mapped to human 22q11 and implicated in schizophrenia. Psychiatr Genet 2015, 25(2):88–94.

25. Sugawara H, Bundo M, Asai T, Sunaga F, Ueda J, Ishigooka J, Kasai K, Kato T, Iwamoto K: Effects of quetiapine on DNA methylation in neuroblastoma cells. Prog Neuropsychopharmacol Biol Psychiatry 2015, 56:117–121.

26. Dong E, Tueting P, Matrisciano F, Grayson DR, Guidotti A: Behavioral and molecular neuroepigenetic alterations in prenatally stressed mice: relevance for the study of chromatin remodeling properties of antipsychotic drugs. Transl Psychiatry 2016, 6:e711.

27. Kinoshita M, Numata S, Tajima A, Yamamori H, Yasuda Y, Fujimoto M, Watanabe S, Umehara H, Shimodera S, Nakazawa T et al: Effect of Clozapine on DNA Methylation in Peripheral Leukocytes from Patients with Treatment-Resistant Schizophrenia. Int J Mol Sci 2017, 18(3).

28. Schizophrenia Working Group of the Psychiatric Genomics C: Biological insights from 108 schizophrenia-associated genetic loci. Nature 2014, 511(7510):421–427.

29. Hannon E, Lunnon K, Schalkwyk L, Mill J: Interindividual methylomic variation across blood, cortex, and cerebellum: implications for epigenetic studies of neurological and neuropsychiatric phenotypes. Epigenetics 2015, 10(11):1024–1032.

30. Bhandari A, Voineskos D, Daskalakis ZJ, Rajji TK, Blumberger DM: A Review of Impaired Neuroplasticity in Schizophrenia Investigated with Non-invasive Brain Stimulation. Front Psychiatry 2016, 7:45.

31. Tiihonen J: Real-world effectiveness of antipsychotics. Acta Psychiatr Scand 2016, 134(5):371–373.

32. Aberg KA, Xie LY, McClay JL, Nerella S, Vunck S, Snider S, Beardsley PM, van den Oord EJ: Testing two models describing how methylome-wide studies in blood are informative for psychiatric conditions. Epigenomics 2013, 5(4):367–377.

33. Rubin LH, Connelly JJ, Reilly JL, Carter CS, Drogos LL, Pournajafi-Nazarloo H, Ruocco AC, Keedy SK, Matthew I, Tandon N et al: Sex and diagnosis specific associations between DNA methylation of the oxytocin receptor gene with emotion processing and temporal-limbic and prefrontal brain volumes in psychotic disorders. Biol Psychiatry Cogn Neurosci Neuroimaging 2016, 1(2):141–151.

34. Saunders JB, Aasland OG, Babor TF, de la Fuente JR, Grant M: Development of the Alcohol Use Disorders Identification Test (AUDIT): WHO Collaborative Project on Early Detection of Persons with Harmful Alcohol Consumption--II. Addiction 1993, 88(6):791–804.

35. Aboraya A, Nasrallah HA: Perspectives on the Positive and Negative Syndrome Scale (PANSS): Use, misuse, drawbacks, and a new alternative for schizophrenia research. Ann Clin Psychiatry 2016, 28(2):125–131.

36. Hu M, Zong X, Zheng J, Mann JJ, Li Z, Pantazatos SP, Li Y, Liao Y, He Y, Zhou J et al: Risperidone-induced topological alterations of anatomical brain network in first-episode drug-naive schizophrenia patients: a longitudinal diffusion tensor imaging study. Psychol Med 2016, 46(12):2549–2560.

37. Hu ML, Zong XF, Zheng JJ, Pantazatos SP, Miller JM, Li ZC, Liao YH, He Y, Zhou J, Sang DE et al: Short-term Effects of Risperidone Monotherapy on Spontaneous Brain Activity in First-episode Treatment-naive Schizophrenia Patients: A Longitudinal fMRI Study. Sci Rep 2016, 6:34287.

38. Jensen AR, Rohwer WD, Jr.: The Stroop color-word test: a review. Acta Psychol (Amst) 1966, 25(1):36–93.

39. Kohli A, Kaur M: Wisconsin Card Sorting Test: Normative data and experience. Indian J Psychiatry 2006, 48(3):181–184.

40. Hazin I, Leite G, Oliveira RM, Alencar JC, Fichman HC, Marques PD, de Mello CB: Brazilian Normative Data on Letter and Category Fluency Tasks: Effects of Gender, Age, and Geopolitical Region. Front Psychol 2016, 7:684.

41. Cheung V, Chiu CP, Law CW, Cheung C, Hui CL, Chan KK, Sham PC, Deng MY, Tai KS, Khong PL et al: Positive symptoms and white matter microstructure in never-medicated first episode schizophrenia. Psychol Med 2011, 41(8):1709–1719.

42. Kates WR, Olszewski AK, Gnirke MH, Kikinis Z, Nelson J, Antshel KM, Fremont W, Radoeva PD, Middleton FA, Shenton ME et al: White matter microstructural abnormalities of the cingulum bundle in youths with 22q11.2 deletion syndrome: associations with medication, neuropsychological function, and prodromal symptoms of psychosis. Schizophr Res 2015, 161(1):76–84.

43. Farkas N, Lendeckel U, Dobrowolny H, Funke S, Steiner J, Keilhoff G, Schmitt A, Bogerts B, Bernstein HG: Reduced density of ADAM 12-immunoreactive oligodendrocytes in the anterior cingulate white matter of patients with schizophrenia. World J Biol Psychiatry 2010, 11(3):556–566.

44. van den Heuvel MP, Mandl RC, Stam CJ, Kahn RS, Hulshoff Pol HE: Aberrant frontal and temporal complex network structure in schizophrenia: a graph theoretical analysis. J Neurosci 2010, 30(47):15915–15926.

45. He Z, Deng W, Li M, Chen Z, Jiang L, Wang Q, Huang C, Collier DA, Gong Q, Ma X et al: Aberrant intrinsic brain activity and cognitive deficit in first-episode treatment-naive patients with schizophrenia. Psychol Med 2013, 43(4):769–780.

46. Howes OD, Kapur S: The dopamine hypothesis of schizophrenia: version III--the final common pathway. Schizophr Bull 2009, 35(3):549–562.

47. Ren W, Lui S, Deng W, Li F, Li M, Huang X, Wang Y, Li T, Sweeney JA, Gong Q: Anatomical and functional brain abnormalities in drug-naive first-episode schizophrenia. Am J Psychiatry 2013, 170(11):1308–1316.

48. Morris TJ, Butcher LM, Feber A, Teschendorff AE, Chakravarthy AR, Wojdacz TK, Beck S: ChAMP: 450k Chip Analysis Methylation Pipeline. Bioinformatics 2014, 30(3):428–430.

49. Teschendorff AE, Marabita F, Lechner M, Bartlett T, Tegner J, Gomez-Cabrero D, Beck S: A beta-mixture quantile normalization method for correcting probe design bias in Illumina Infinium 450 k DNA methylation data. Bioinformatics 2013, 29(2):189–196.

50. Nordlund J, Backlin CL, Wahlberg P, Busche S, Berglund EC, Eloranta ML, Flaegstad T, Forestier E, Frost BM, Harila-Saari A et al: Genome-wide signatures of differential DNA methylation in pediatric acute lymphoblastic leukemia. Genome Biol 2013, 14(9):r105.

51. Johnson WE, Li C, Rabinovic A: Adjusting batch effects in microarray expression data using empirical Bayes methods. Biostatistics 2007, 8(1):118–127.

52. Horvath S: DNA methylation age of human tissues and cell types. Genome Biol 2013, 14(10):R115.

53. Saffari A, Silver MJ, Zavattari P, Moi L, Columbano A, Meaburn EL, Dudbridge F: Estimation of a significance threshold for epigenome-wide association studies. Genet Epidemiol 2018, 42(1):20–33.

54. Ritchie ME, Phipson B, Wu D, Hu Y, Law CW, Shi W, Smyth GK: limma powers differential expression analyses for RNA-sequencing and microarray studies. Nucleic acids research 2015, 43(7):e47.

55. Martorell-Marugan J, Gonzalez-Rumayor V, Carmona-Saez P: mCSEA: Detecting subtle differentially methylated regions. Bioinformatics 2019.

56. Wu Y, Yao YG, Luo XJ: SZDB: A Database for Schizophrenia Genetic Research. Schizophr Bull 2017, 43(2):459–471.

57. Li J, Cai T, Jiang Y, Chen H, He X, Chen C, Li X, Shao Q, Ran X, Li Z et al: Genes with de novo mutations are shared by four neuropsychiatric disorders discovered from NPdenovo database. Mol Psychiatry 2016, 21(2):290–297.

58. Phipson B, Maksimovic J, Oshlack A: missMethyl: an R package for analyzing data from Illumina's HumanMethylation450 platform. Bioinformatics 2016, 32(2):286–288.

59. Bishop JR, Reilly JL, Harris MS, Patel SR, Kittles R, Badner JA, Prasad KM, Nimgaonkar VL, Keshavan MS, Sweeney JA: Pharmacogenetic associations of the type-3 metabotropic glutamate receptor (GRM3) gene with working memory and clinical symptom response to antipsychotics in first-episode schizophrenia. Psychopharmacology (Berl) 2015, 232(1):145–154.

60. Shannon P, Markiel A, Ozier O, Baliga NS, Wang JT, Ramage D, Amin N, Schwikowski B, Ideker T: Cytoscape: a software environment for integrated models of biomolecular interaction networks. Genome Res 2003, 13(11):2498–2504.

