## Supplementary Files for "Risperidone-induced changes in DNA methylation from peripheral blood in first-episode schizophrenia parallel neuroimaging and cognitive phenotype"

### Supplementary material

**Figure S1. Overall methylation-phenotype network.**

Pink line means the correlation between the two nodes is negative, grey line means the correlation is positive. The width of the line represents the absolute value of correlation coefficient.

**Figure S2. Methylation-phenotype network for PANSS scores.**

**Figure S3. Methylation-phenotype network for fMRI.**

Table S1. description of brain function variables and replication data. (A) brain activity variables; (B) brain topological network variables; (C) cognitive function variables; (D) replication data set.

Table S2. Normalized genes overlapped with schizophrenia related genes.

Table S3. Correlation of methylation in calcium genes with brain function variables.

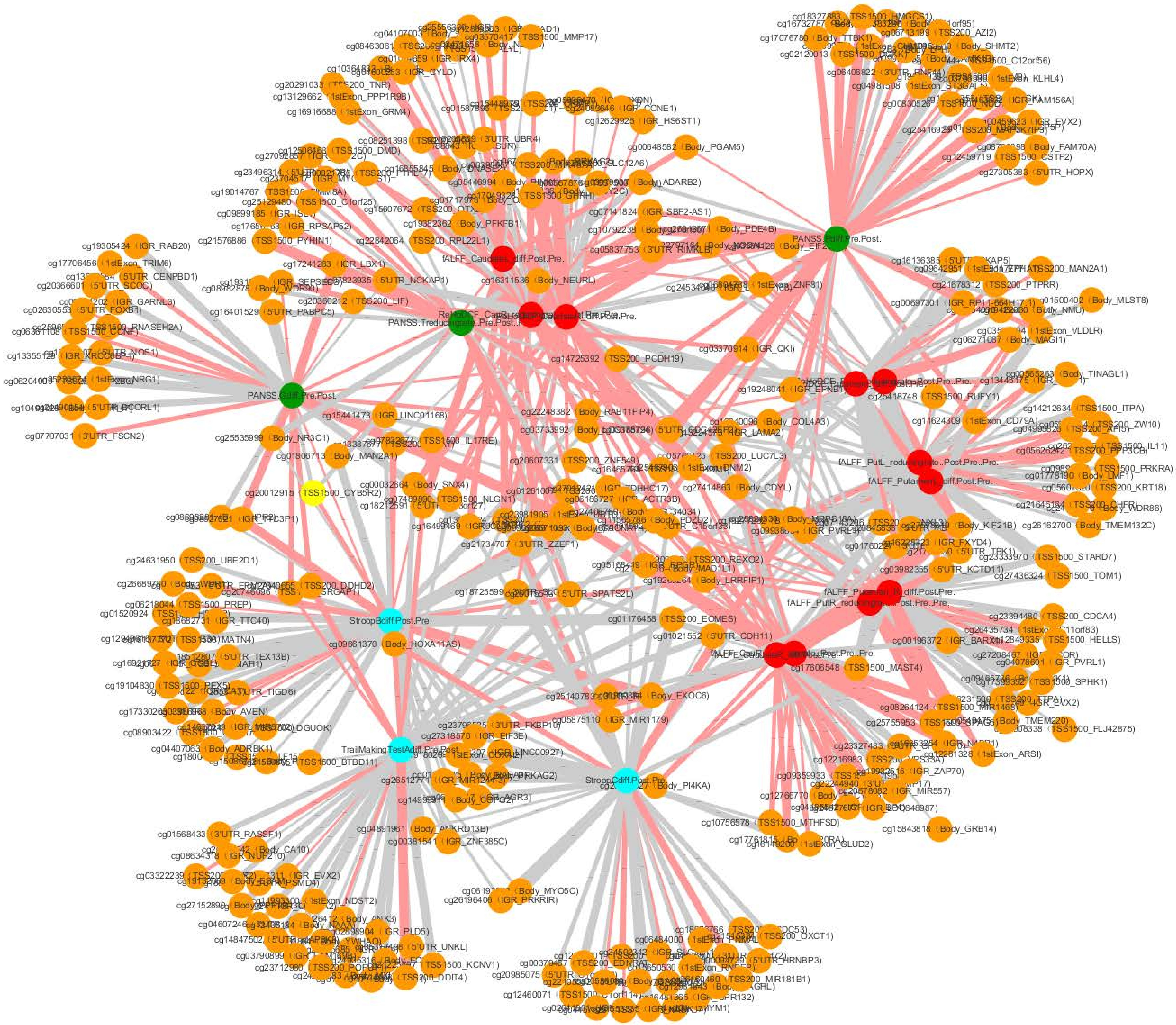
PANSS.Pdiff.Pre.Post.
StroopBdiff.Post.Pre.
StroopCdiff.Post.Pre.
cg19305424 IGR_RAB20
cg17706456 1stExon_TRIM6
cg07707031 3'UTR_FSCN2
cg24631950 TSS200_UBE2D1
cg01568433 3'UTR_RASSF1
cg15843818 Body_GRB14
cg27305383 5'UTR_HOPX
cg12459719 TSS1500_CSTF2
cg00648582 Body_PGAM5
cg12629925 IGR_HS6ST1
cg14212634 TSS1500_ITPA
cg23394480 TSS200_CDCA4
cg10756578 TSS1500_MTHFSD
cg01176458 TSS200_EOMES
cg14725392 TSS200_PCDH19
cg03370914 IGR_QKI
cg26162700 Body_TMEM132C
cg27436324 TSS1500_TOM1
cg24891961 Body_ANKRD13B
cg00381541 IGR_ZNF385C
cg16311536 Body_NEURL
cg21576886 TSS1500_PYHIN1
cg09661370 Body_HOXA11AS
cg17241283 IGR_LBX1
cg01021552 5'UTR_CDH11
cg21734707 3'UTR_ZZEF1
cg16401529 5'UTR_PABPC5
cg25535999 Body_NR3C1
cg21678312 TSS200_PTPRR
cg06271087 Body_MAGI1

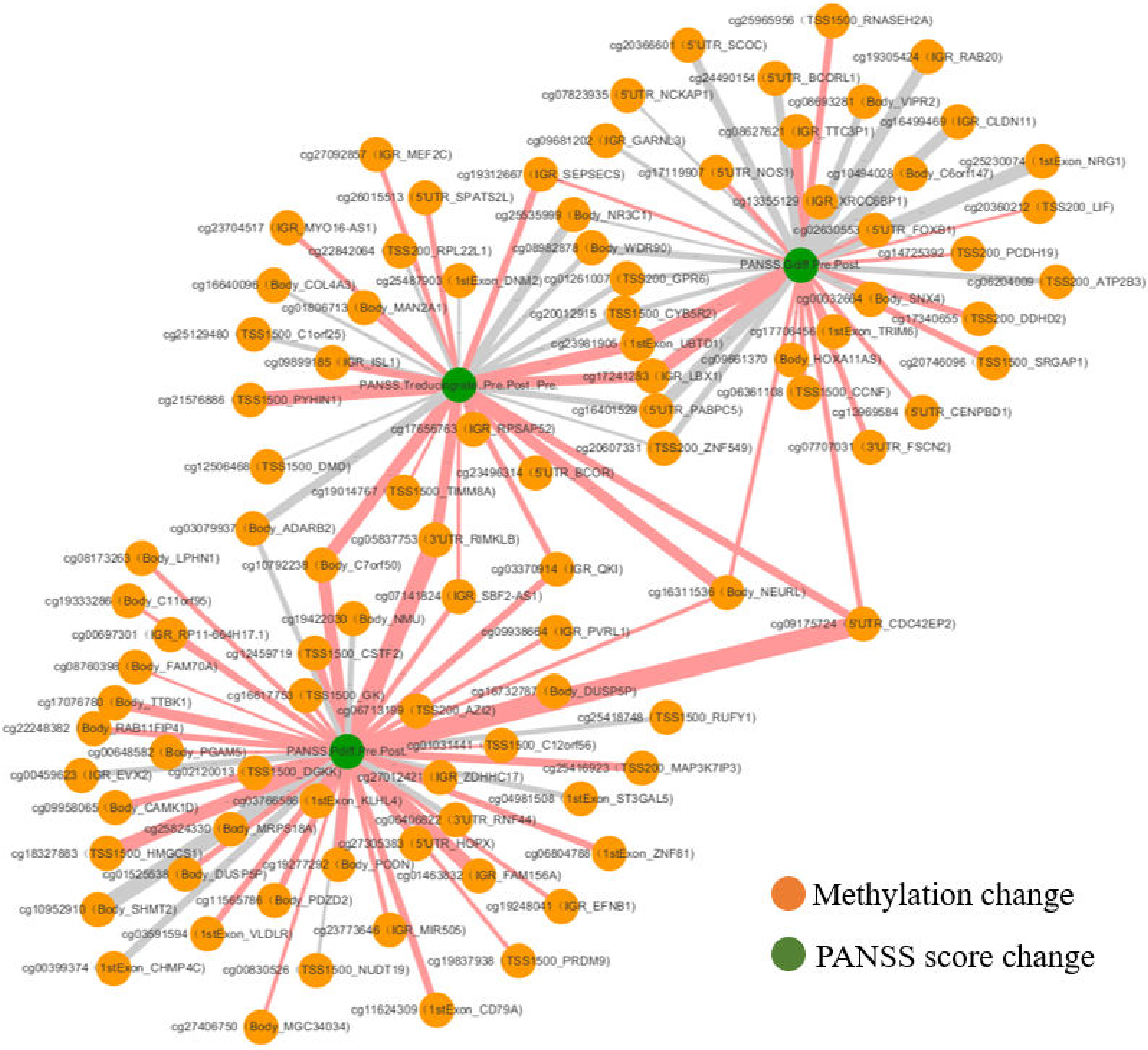
cg25965956（TSS1500_RNASEH2A)
cg20366601（5'UTR_SCOC)
cg19305424（IGR_RAB20)
cg24490154（5'UTR_BCORL1)
cg07823935（5'UTR_NCKAP1)
cg08693281（Body_VIPR2)
cg16499469（IGR_CLDN11)
cg08627621（IGR_TTC3P1)
cg09681202（IGR_GARNL3)
cg27092857（IGR_MEF2C)
cg25230074（1stExon_NRG1)
cg19312667（IGR_SEPSECS)
cg17119907（5'UTR_NOS1)
cg10484028（Body_C6orf147)
cg26015513（5'UTR_SPATS2L)
cg13355129（IGR_XRCC6BP1)
cg20360212（TSS200_LIF)
cg25535999（Body_NR3C1)
cg23704517（IGR_MYO16-AS1)
cg02630553（5'UTR_FOXB1)
cg08982878（Body_WDR90)
cg22842064（TSS200_RPL22L1)
cg14725392（TSS200_PCDH19)
PANSS.Gdiff.Pre.Post.
cg25487903（1stExon_DNM2)
cg01261007（TSS200_GPR6)
cg16640096（Body_COL4A3)
cg06204009（TSS200_ATP2B3)
cg01806713（Body_MAN2A1)
cg00032664（Body_SNX4)
cg20012915（TSS1500_CYB5R2)
cg17340655（TSS200_DDHD2)
cg25129480（TSS1500_C1orf25)
cg17706456（1stExon_TRIM6)
cg23981906（1stExon_UBTD1)
cg09899185（IGR_ISL1)
cg09661370（Body_HOXA11AS)
cg20746096（TSS1500_SRGAP1)
cg17241283（IGR_LBX1)
cg21576886（TSS1500_PYHIN1)
cg06361108（TSS1500_CCNF)
cg16401529（5'UTR_PABPC5)
cg13969584（5'UTR_CENPBD1)
cg17656763（IGR_RPSAP52)
cg20607331（TSS200_ZNF549)
cg07707031（3'UTR_FSCN2)
cg12506468（TSS1500_DMD)
cg23496314（5'UTR_BCOR)
cg19014767（TSS1500_TIMM8A)
cg03079937（Body_ADARB2)
cg05837753（3'UTR_RIMKLB)
cg08173263（Body_LPHN1)
cg10792238（Body_C7orf50)
cg03370914（IGR_QKI)
cg07141824（IGR_SBF2-AS1)
cg16311536（Body_NEURL)
cg19333286（Body_C11orf95)
cg19422030（Body_NMU)
cg09175724（5'UTR_CDC42EP2)
cg00697301（IGR_RP11-664H17.1)
cg09938664（IGR_PVRL1)
cg12459719（TSS1500_CSTF2)
cg08760398（Body_FAM70A)
cg16732787（Body_DUSP5P)
cg17076780（Body_TTBK1)
cg16817753（TSS1500_GK)
cg06713199（TSS200_AZI2)
cg25418748（TSS1500_RUFY1)
cg22248382（Body_RAB11FIP4)
cg00648582（Body_PGAM5)
PANSS.Pdiff.Pre.Post.
cg01031441（TSS1500_C12orf56)
cg00459623（IGR_EVX2)
cg02120013（TSS1500_DGKK)
cg25416923（TSS200_MAP3K7IP3)
cg27012421（IGR_ZDHHC17)
cg09958065（Body_CAMK1D)
cg03766586（1stExon_KLHL4)
cg04981508（1stExon_ST3GAL5)
cg25824330（Body_MRPS18A)
cg08406822（3'UTR_RNF44)
cg18327883（TSS1500_HMGCS1)
cg27305383（5'UTR_HOPX)
cg06804788（1stExon_ZNF81)
cg01525538（Body_DUSP5P)
cg19277292（Body_PODN)
cg01463832（IGR_FAM156A)
cg10952910（Body_SHMT2)
cg11565786（Body_PDZD2)
cg19248041（IGR_EFNB1)
cg03591594（1stExon_VLDLR)
cg23773646（IGR_MIR505)
cg00399374（1stExon_CHMP4C)
cg00830526（TSS1500_NUDT19)
cg19837938（TSS1500_PRDM9)
cg11624309（1stExon_CD79A)
cg27406750（Body_MGC34034)
Methylation change
PANSS score change

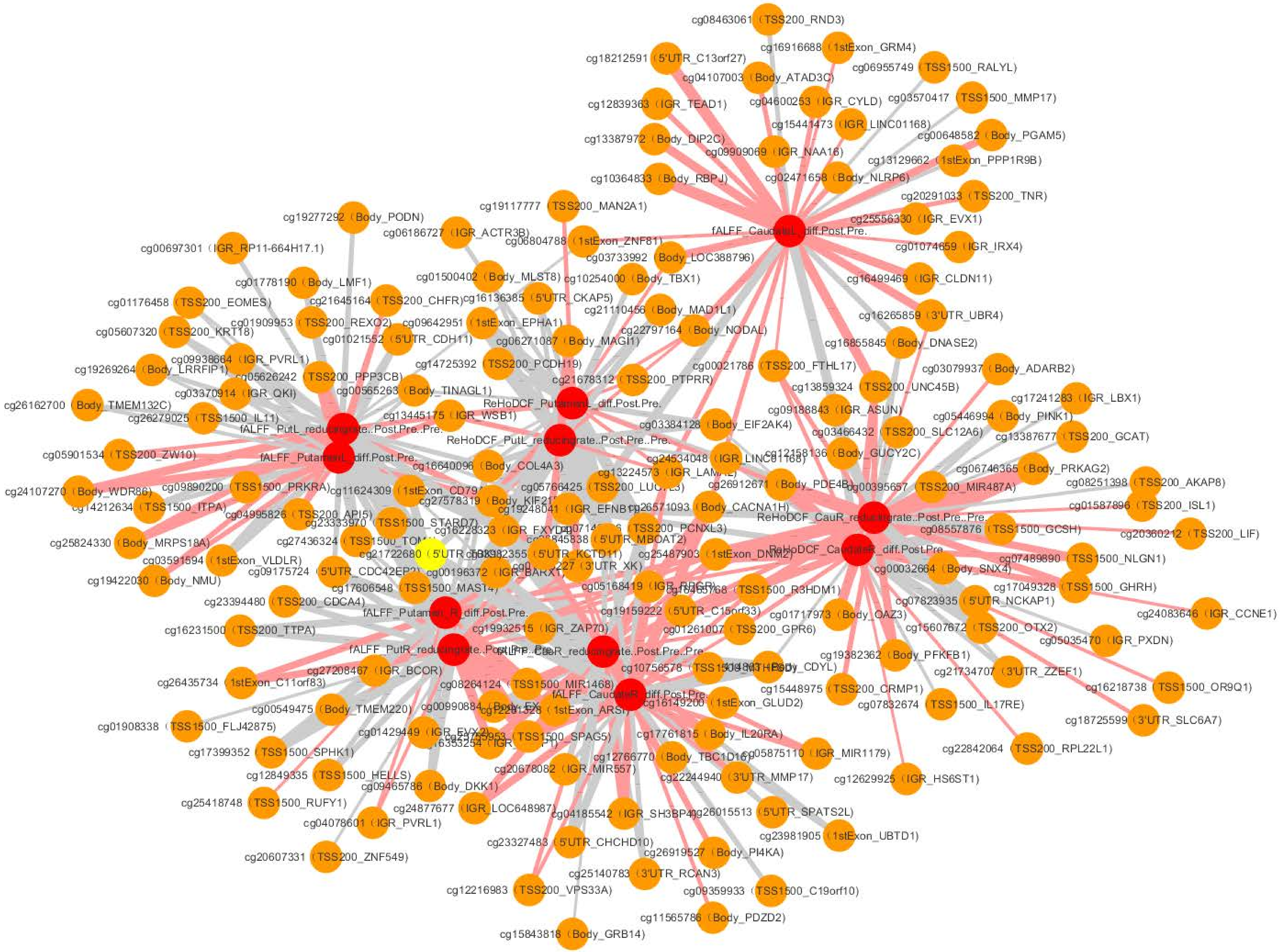
cg08463061（TSS200_RND3)
cg16916688（1stExon_GRM4)
cg18212591（5'UTR_C13orf27)
cg06955749（TSS1500_RALYL)
cg04107003（Body_ATAD3C)
cg04600253（IGR_CYLD)
cg03570417（TSS1500_MMP17)
cg12839363（IGR_TEAD1)
cg15441473（IGR_LINC01168)
cg13387972（Body_DIP2C)
cg00648582（Body_PGAM5)
cg09909069（IGR_NAA16)
cg13129662（1stExon_PPP1R9B)
cg10364833（Body_RBPJ)
cg02471658（Body_NLRP6)
cg20291033（TSS200_TNR)
cg19117777（TSS200_MAN2A1)
cg25556330（IGR_EVX1)
fALFF_CaudateL_diff.Post.Pre.
cg19277292（Body_PODN)
cg06186727（IGR_ACTR3B)
cg06804788（1stExon_ZNF81)
cg01074659（IGR_IRX4)
cg00697301（IGR_RP11-664H17.1)
cg03733992（Body_LOC388796)
cg01778190（Body_LMF1)
cg01500402（Body_MLST8)
cg10254000（Body_TBX1)
cg16499469（IGR_CLDN11)
cg01176458（TSS200_EOMES)
cg21645164（TSS200_CHFR)
cg16136385（5'UTR_CKAP5)
cg21110456（Body_MAD1L1)
cg16265859（3'UTR_UBR4)
cg01909953（TSS200_REXO2)
cg09642951（1stExon_EPHA1)
cg05607320（TSS200_KRT18)
cg01021552（5'UTR_CDH11)
cg06271087（Body_MAGI1)
cg22797164（Body_NODAL)
cg16855845（Body_DNASE2)
cg09938664（IGR_PVRL1)
cg14725392（TSS200_PCDH19)
cg00021786（TSS200_FTHL17)
cg03079937（Body_ADARB2)
cg19269264（Body_LRRFIP1)
cg05626242（TSS200_PPP3CB)
cg21678312（TSS200_PTPRR)
cg13859324（TSS200_UNC45B)
cg03370914（IGR_QKI)
cg00565263（Body_TINAGL1)
cg17241283（IGR_LBX1)
cg26162700（Body_TMEM132C)
ReHoDCF_PutamenL_diff.Post.Pre.
cg09188843（IGR_ASUN)
cg05446994（Body_PINK1)
cg26279025（TSS1500_IL11)
cg13445175（IGR_WSB1)
cg03384128（Body_EIF2AK4)
cg03466432（TSS200_SLC12A6)
cg13387677（TSS200_GCAT)
fALFF_PutL_reducingrate..Post.Pre..Pre.
ReHoDCF_PutL_reducingrate..Post.Pre..Pre.
cg05901534（TSS200_ZW10)
fALFF_PutamenL_diff.Post.Pre.
cg12158136（Body_GUCY2C)
cg06746365（Body_PRKAG2)
cg16640096（Body_COL4A3)
cg08251398（TSS200_AKAP8)
cg24107270（Body_WDR86)
cg09890200（TSS1500_PRKRA)
cg26912671（Body_PDE4B)
cg00395657（TSS200_MIR487A)
cg14212634（TSS1500_ITPA)
cg26571093（Body_CACNA1H)
cg01587896（TSS200_ISL1)
cg04995826（TSS200_API5)
ReHoDCF_CauR_reducingrate..Post.Pre..Pre.
cg25824330（Body_MRPS18A)
cg08557876（TSS1500_GCSH)
cg20360212（TSS200_LIF)
cg25487903（1stExon_DNM2)
ReHoDCF_CaudateR_diff.Post.Pre.
cg03591594（1stExon_VLDLR)
cg07489690（TSS1500_NLGN1)
cg19422030（Body_NMU)
cg00032664（Body_SNX4)
cg17049328（TSS1500_GHRH)
cg23394480（TSS200_CDCA4)
cg07823935（5'UTR_NCKAP1)
fALFF_PutamenR_diff.Post.Pre.
cg24083646（IGR_CCNE1)
cg16231500（TSS200_TTPA)
cg19932515（IGR_ZAP70)
cg15607672（TSS200_OTX2)
cg05035470（IGR_PXDN)
fALFF_PutR_reducingrate..Post.Pre..Pre.
fALFF_CauR_reducingrate..Post.Pre..Pre.
cg19382362（Body_PFKFB1)
cg27208467（IGR_BCOR)
cg21734707（3'UTR_ZZEF1)
cg26435734（1stExon_C11orf83)
cg08264124（TSS1500_MIR1468)
cg15448975（TSS200_CRMP1)
cg16218738（TSS1500_OR9Q1)
fALFF_CaudateR_diff.Post.Pre.
cg00549475（Body_TMEM220)
cg16149200（1stExon_GLUD2)
cg07832674（TSS1500_IL17RE)
cg18725599（3'UTR_SLC6A7)
cg01908338（TSS1500_FLJ42875)
cg17761815（Body_IL20RA)
cg17399352（TSS1500_SPHK1)
cg05875110（IGR_MIR1179)
cg22842064（TSS200_RPL22L1)
cg12849335（TSS1500_HELLS)
cg12766770（Body_TBC1D16)
cg09465786（Body_DKK1)
cg20678082（IGR_MIR557)
cg22244940（3'UTR_MMP17)
cg12629925（IGR_HS6ST1)
cg25418748（TSS1500_RUFY1)
cg24877677（IGR_LOC648987)
cg04078601（IGR_PVRL1)
cg04185542（IGR_SH3BP4)
cg26015513（5'UTR_SPATS2L)
cg23981905（1stExon_UBTD1)
cg23327483（5'UTR_CHCHD10)
cg26919527（Body_PI4KA)
cg20607331（TSS200_ZNF549)
cg25140783（3'UTR_RCAN3)
cg12216983（TSS200_VPS33A)
cg09359933（TSS1500_C19orf10)
cg11565786（Body_PDZD2)
cg15843818（Body_GRB14)

**Supplementary Table S1A.** Baseline and longitudinal alterations in fALFF and ReHo in striatum ($p<0.05$, AlphaSim corrected)

| Brain region | AAL | MNI coordinates | | | Voxels | Maximat *t* value |
|---|---|---|---|---|---|---|
| | | X | Y | Z | | |
| Patient at baseline *vs* Controls | | | | | | |
| fALFF, left caudate | 71 | -15 | 3 | 18 | 52 | 5.96 |
| Patient at follow-up vs Baseline | | | | | | |
| fALFF, right caudate | 72 | 12 | 15 | 0 | 71 | 5.43 |
| fALFF, left putamen | 73 | -21 | 15 | 3 | 38 | 5.24 |
| fALFF, right putamen | 74 | 21 | 18 | 0 | 50 | 5.39 |
| ReHo, right caudate | 72 | 12 | 12 | -9 | 20 | 4.75 |
| ReHo, left putamen | 73 | -21 | 12 | 12 | 20 | 5.12 |

AAL= Automated Anatomical Labeling; MNI= Montreal Neurological Institute; fALFF= fractional amplitude of low-frequency fluctuation; ReHo= regional homogeneity. Multiple correction was performed using cluster-extent correction (AlphaSim) as follows: individual voxel threshold p=0.001, Number of Monte Carlo simulations=1000, and p=0.05 as the effective threshold for cluster-extent correction.

**Supplementary Table S1B.** Nodal topological characteristics of anatomical brain networks in healthy controls and first-episode schizophrenia patients for baseline and follow-up data (FDR corrected)

| Nodal Property | Controls | Patients at baseline | Patients at follow-up | ***P* value** | | |
|---|---|---|---|---|---|---|
| | | | | **patients at baseline *vs.* controls** | **patients at baseline *vs.* follow-up** | **patients at follow-up *vs.* controls** |
| Left ACG/AAL31 | | | | | | |
| $k_i^W$ | 1022.0±836.0 | 671.8±514.4 | 868.2±824.1 | 0.035 | 0.043 | 0.066 |
| Right ACG/AAL32 | | | | | | |
| $k_i^W$ | 702.0±740.0 | 423.6±453.7 | 555.7±596.7 | 0.032 | 0.023 | 0.062 |
| $E_{nodal}$ | 45.5±24.8 | 33.8±20.7 | 38.3±23.1 | 0.040 | 0.048 | 0.076 |
| Left PCG/AAL35 | | | | | | |
| $CC_{nodal}$ | 54.9±24.5 | 43.6±18.4 | 48.6±21.7 | 0.035 | 0.042 | 0.056 |
| Right PCG/AAL36 | | | | | | |
| $CC_{nodal}$ | 54.3±21.4 | 45.1±17.3 | 49.6±22.8 | 0.048 | 0.049 | 0.060 |
| Left SFG_med_orb/AAL25 | | | | | | |
| $k_i^W$ | 459.9±332.4 | 245.6±179.7 | 292.7±213.7 | 0.019 | 0.052 | 0.019 |
| $E_{nodal}$ | 32.3±15.1 | 22.1±10.3 | 24.7±11.2 | 0.013 | 0.079 | 0.041 |
| Right pallidum/AAL76 | | | | | | |
| $k_i^W$ | 937.9±555.6 | 721.0±342.4 | 815.8±416.4 | 0.020 | 0.060 | 0.049 |
| left amygdala/AAL41 | | | | | | |
| $k_i^W$ | 150.7±138.2 | 115.7±128.1 | 172.7±128.6 | 0.056 | 0.003 | 0.071 |
| $E_{nodal}$ | 18.1±10.8 | 15.7±8.7 | 20.4±8.5 | 0.071 | 0.005 | 0.055 |

| | | | | | | |
|---|---|---|---|---|---|---|
| left PHG/AAL39 | | | | | | |
| $k_i^W$ | 138.1±100.6 | 175.4±153.6 | 182.8±111.8 | 0.053 | 0.044 | 0.035 |
| $E_{nodal}$ | 16.7±8.8 | 17.7±8.6 | 20.2±9.7 | 0.078 | 0.030 | 0.053 |
| left CAU/AAL71 | | | | | | |
| $k_i^W$ | 1065.6±489.3 | 892.6±427.1 | 769.4±387.6 | 0.052 | 0.013 | 0.015 |

*FDR = False Discovery Rate; AAL = Automated Anatomical Labeling atlas; ACG = anterior cingulate and paracingulate gyri; PCG=Posterior cingulate gyrus; SFG_med_orb = superior frontal gyrus medial orbital; PHG = parahippocampal gyrus; CAU = caudate nucleus;* $k_i^W$ *= nodal degree; CCnodal =clustering coefficiency;* $E_{nodal}$ *= nodal efficiency.*

**Supplementary Table S1C.** Cognitive function in healthy controls and first-episode schizophrenia patients for baseline and follow-up data

| cognitive function | Patients（n=38） | | Controls（n=38） | Pre-treatment vs control | | Pre-treatment vs Post-treatment | | Post-treatment vs control | |
|---|---|---|---|---|---|---|---|---|---|
| | Pre-treatment | Post-treatment | | t | p | t | P | t | p |
| WCST-PE | 50.26±6.96 | 47.87±9.83 | 32.0 ± 10.35 | 9.03 | <0.001 | 1.298 | 0.202 | 6.85 | <0.001 |
| WCST-categories | 3.92±0.82 | 3.47±1.08 | 4.53±1.06 | -2.79 | 0.007 | -1.769 | 0.077 | -4.28 | 0.001 |
| DSDT-forward | 7.5±1.2 | 7.7±1.3 | 8.7±1.3 | -4.07 | <0.001 | 0.79 | 0.438 | -3.41 | 0.001 |
| DSDT-backward | 4.9±1.4 | 4.6±1.1 | 6.5±1.4 | -4.80 | <0.001 | 1.82 | 0.076 | -6.28 | <0.001 |
| VFT | 17.0±5.9 | 17.1±5.0 | 23.2±5.5 | -4.74 | <0.001 | 0.13 | 0.899 | -5.08 | <0.001 |
| SCWT-A | 23.9±12.0 | 22.1±9.1 | 14.7±4.1 | 4.45 | <0.001 | 1.23 | 0.228 | 4.55 | <0.001 |
| SCWT-B | 27.0±12.4 | 24.3±9.6 | 15.9±4.3 | -5.20 | <0.001 | 2.38 | 0.022 | 4.91 | <0.001 |
| SCWT-C | 39.7±18.5 | 35.3±12.1 | 27.2±8.6 | 3.77 | <0.001 | 2.22 | 0.033 | 3.36 | 0.001 |
| TMT-A | 63.2±39.8 | 51.5±28.5 | 34.7±13.5 | 4.18 | <0.001 | 2.03 | 0.05 | 3.28 | 0.002 |
| TMT-B | 162.0±84.6 | 135.6±82.8 | 71.8±21.8 | 5.77 | <0.001 | 1.89 | 0.067 | 4.59 | <0.001 |

Wisconsin Card Sorting Test (WCST), Stroop Color Word Test (SCWT), Trail Making Test (TMT), Verbal Fluency Test (VFT), Digit Span Distraction Test (DSDT). Persistence errors (PE)

**Supplementary Table S1D.** **General information of the replication data**

| Individuals | **Sex** | **Race** | **Profile** | **Age** |
|---|---|---|---|---|
| FEP-106 | Male | African Am | Control | 30 |
| FEP-109 | Male | African Am | Control | 27 |
| FEP-105 | Female | Hispanic | Control | 28 |
| FEP-125 | Female | White | Control | 27 |
| FEP-107 | Female | African Am | Schizophrenia | 22 |
| FEP-108 | Female | Hispanic | Schizophrenia | 40 |
| FEP-113 | Male | African Am | Schizophrenia | 18 |
